# Decoupled calcium homeostasis and signaling associated with cytoskeletal instability in YWHAG R132C induced pluripotent stem cell-derived cortical neurons

**DOI:** 10.64898/2026.04.01.715876

**Authors:** Anna Maria Schreiber, Anamika Gupta, Ashley Thompson, Dev Raj Bhattarai, Russell D’Souza, Lindsay Rizzardi, João Pereira

## Abstract

YWHAG Syndrome (Developmental and Epileptic Encephalopathy 56, DEE56) is an ultra- rare childhood epilepsy associated with neurodevelopmental delays, with no therapeutic intervention available. Multiple *de novo* mutations in the *YWHAG* gene, encoding for the 14-3-3γ protein, have been identified as causative for YWHAG Syndrome. 14-3-3γ interacts with various targets, including major neurodevelopmental signaling proteins such as components of the ROCK pathway. Despite substantial evidence of the essential role of 14-3-3γ in neurite outgrowth, cytoskeletal rearrangements, and neuronal migration during cortical development, little is known regarding the molecular consequences of *YWHAG* mutations and their effect on neuronal function and survival. Here, we characterized an isogenic, pluripotent stem cell (iPSC) model of *YWHAG^R132C/+^* cortical neurons. The *YWHAG^R132C/+^*iPSC-derived neurons exhibited early cytoskeletal phenotypes, coupled with an elevated calcium baseline, lower frequency of calcium spikes, and reduced network activity. The widespread alterations in the transcriptome of mutant neurons revealed a biphasic dysregulation in the core genes and modulators associated with the ROCK pathway that resulted in maturation-dependent changes to cytoskeletal protein stability and calcium phenotypes. Direct inhibition of ROCK with Y27632 further increased the calcium baseline compared to the isogenic control. Exposure of *YWHAG*^R132C/+^ neurons to Trypsin-EDTA revealed underlying cytoskeletal instability, which was partially reversed by lovastatin treatment. Further, lovastatin partially rescued the elevated calcium baseline, but not the frequency or amplitude of calcium spikes. Together, these results suggest decoupling of calcium homeostasis and calcium signaling associated with cytoskeletal instability in *YWHAG*^R132C/+^ neurons. These findings lay the groundwork for future mechanistic studies of YWHAG function and molecular therapeutic targets for YWHAG Syndrome and YWHAG-associated conditions.

## INTRODUCTION

YWHAG Syndrome, or Developmental and Epileptic Encephalopathy 56 (DEE56; OMIM #617665), is an ultra-rare genetic condition presenting with early-onset epilepsy, including epileptic encephalopathy (EE) and drug-resistant seizures, and accompanied by neurodevelopmental delays^1–5^. The Epi4K Consortium & Epilepsy Phenome/Genome Project study confirmed that *de novo* mutations in the *YWHAG* gene, encoding for the 14-3-3γ protein, are causative for YWHAG Syndrome^1^. Critically, little is known about the molecular mechanisms underlying YWHAG Syndrome, and no therapeutic interventions are available to patients.

The 14-3-3γ protein (Tyrosine 3-Monooxygenase/Tryptophan 5-Monooxygenase Activation Protein Gamma) is a member of the highly conserved 14-3-3 family of proteins. The 14-3-3s contribute to various biological functions, including signal transduction mediation through binding to phosphorylated serine and threonine motifs on target peptides^6^. 14-3-3 dimers bind target proteins at a structural U-shaped groove consisting of nine alpha-helical structures^7^. Expression of the 14-3-3 protein family in the brain is ubiquitous, with isoforms varying across cell types and developmental stages^8^. 14-3-3γ, encoded by *YWHAG*, is one of the predominant neuronal isoforms of 14-3-3 during development^9^. 14-3-3γ is highly expressed during the embryonic stage in mice until P7, when expression gradually decreases, reaching a minimal level by P30^10^. 14-3-3γ plays a crucial role in neuronal migration during this window of early cortical development, and *Ywhag* ablation results in delays in neuronal migration and morphological defects^10,11^. YWHAG also interacts with major neurodevelopmental signaling proteins, including components of the ROCK pathway, RAF1, and Protein Kinase C (PKC)^12,13^, suggesting a role in neurite outgrowth and cytoskeletal rearrangements. Multiple missense point mutations in *YWHAG* have been identified as disease-causing, with the most severe located within the binding groove of 14-3-3γ^4,5^.

Here, we set out to address the molecular and cellular phenotypes resulting from the *YWHAG^R132C^* mutation using an isogenic model of human induced pluripotent stem cells (iPSCs). We differentiated and characterized NGN2-derived cortical excitatory neurons, profiling changes to cytoskeletal stability, transcriptomics, calcium homeostasis, calcium event amplitude and frequency, and network excitability. We uncovered a *YWHAG^R132C^*neuronal phenotype of cytoskeletal instability and an elevated baseline calcium level, associated with reduced spike frequency and amplitude. We found that the ROCK pathway modulates a biphasic response to *YWHAG^R132C^*, first increasing and later suppressing the expression of focal actin genes. Critically, treatment with lovastatin stabilized the cytoskeleton and decoupled the calcium baseline from calcium spike amplitude and frequency in *YWHAG^R132C/+^* ^neurons^, supporting a link between cytoskeletal and calcium homeostasis. The characterization of cellular and molecular *YWHAG^R132C^* phenotypes lays the groundwork for future mechanistic studies of YWHAG function and YWHAG-associated conditions.

## METHODS

### Induced pluripotent stem cell culture and maintenance

This study utilized the iP11NK induced pluripotent stem cell (iPSC) line (ALSTEM), edited to insert the *YWHAG* R132C mutation (Transcripta Bio), matched with the isogenic control line. Both lines have a Tet-ON::NGN2 (Neurogenin) expression cassette inserted into the AAVS1 safe harbor locus. Cells were grown in Matrigel^®^ (Corning, 354277) coated 6-well plates (Fisher brand, FB012927), in mTeSR Plus medium (STEMCELL Technologies, 100-0276) supplemented with 10 μM Y-27632 Rho kinase inhibitor (abcam, ab144494) with daily medium changes.

### Differentiation of doxycycline-inducible NGN2 cortical neurons

iPSCs were dissociated using StemPro™ Accutase™ Cell Dissociation Reagent (Gibco™, A11105), and 4 million cells were plated on Matrigel®-coated (Corning, 354277) 10-centimeter dishes (Thermo Fisher Scientific, 130182) in mTeSRPlus, supplemented with 10 μM Y-27632 Rho kinase inhibitor. For the following three days, mTeSR Plus was replaced with neuronal pre-differentiation medium containing Knockout DMEM/F12 (Gibco™, 12660012), supplemented with 1X MEM Non-Essential Amino Acids (Gibco™,11140050), 1X N2 Supplement (Gibco™, 17502048), 10 ng/mL NT-3 (PeproTech, 450-03), 10 ng/mL BDNF (PeproTech, 450-02), 1 ug/mL Mouse Laminin (Gibco™, 23017015), 10 μM Y-27632 Rho kinase inhibitor, and 2 ug/mL doxycycline-hydrochloride (Sigma-Aldrich, D3072). Following three days of neuronal pre-differentiation, cells were passaged onto poly-L-lysine (EMD Milipore, P4707)/laminin (Gibco™, 23017015)-coated plates at densities of 25,000 cells/well for 24-well plates (Cellvis, P24-1.5H-N), and 4 million cells for 10-centimeter dishes. Cells were maintained in classic neuronal medium consisting of 1:1 ratio of DMEM/F12 (Gibco™, 10565018): Neurobasal-A (Gibco™, 10888022), 1X MEM Non-Essential Amino Acids, 0.5X GlutaMAX Supplement (Gibco™, 35050061), 0.5X N2 Supplement, 0.5X B27 Supplement (Gibco™, 17504044), 10 ng/mL NT-3, 10 ng/mL BDNF, 1 ug/mL Mouse Laminin. Half of the medium change was performed every 7 days^14^.

### Immunocytochemistry

Differentiated cortical neurons were fixed using 4% paraformaldehyde (v/v) (Electron Microscopy Sciences, 15710) in PBS (Gibco™,100-10-023), for 20 minutes at room temperature, followed by 1-hour of primary antibody incubation diluted in PBS with 0.1% of Triton-X (Thermo Fisher Scientific, A16046-0F). Following a PBS wash, cells were incubated with Alexa Fluor™-labeled secondary antibodies diluted in PBS (1:1000) with 0.1% Triton-X and incubated for 1 hour. Nuclei were then labeled with DAPI (Thermo Fisher Scientific, D9542) (1:10,000), and cells were washed with PBS. The following primary antibodies and dilutions were used: anti-SOX2 (1:500, abcam, ab97959), anti-Caspase3 (1:500, Cell Signaling Technologies, 9661S), anti-MAP2 (1:500, abcam, ab32454), anti-NeuN (1:500, Milipore Sigma, MAB377X), anti-14-3-3γ (1:500, GeneTex, GTX113298). The following secondary antibodies were used: Alexa Fluor™ 488 anti-mouse (Invitrogen, A21202), Alexa Fluor™ 488 anti-chicken (Invitrogen, A78948), Alexa Fluor™ 647 anti-rabbit (Invitrogen, A31573), Alexa Fluor™ 647 anti-mouse (Invitrogen, A31571). All images were acquired using the ImageXpress® Confocal HT.ai High-Content Imaging System (Molecular Devices, LLC).

### Calcium imaging

Neuronal cultures were loaded with 5 μM Fluo-4 AM (Thermo Fisher Scientific, F14201), a cell-permeant dye, reconstituted in DMSO (Sigma-Aldrich, D2650) according to the manufacturer’s instructions. After 30 minutes of incubation, epifluorescence was acquired using the ImageXpress® HT.ai. Acquisition settings were defined with 4x binning, 30 ms exposure, and 10% power for the 488 laser, with care to maintain the initial signal within half of the camera’s dynamic range. Autofocus was enabled for all sites, and 30-second movies were acquired at 30 FPS per site, for a total of 8 sites per well. Downstream analyses were conducted in FIJI^15^ using custom macros. Briefly, all individual frames of each 30-second movie (900 frames/movie) were opened as a stack, and an average intensity projection was generated. The average projection image was converted to 8-bit, the background was subtracted, and the resulting image was automatically thresholded. A mask of particles within the 50-600 px^2^ range was generated and transferred as ROIs (regions of interest) to the ROI manager, enabling quantification of Fluo-4 AM intensity per image in the stack and over time, as well as downstream statistical analysis.

### Multielectrode array recordings

Following the three days of neuronal pre-differentiation as described above, the cells were plated onto a poly-L-lysine/laminin-coated CytoView MEA 48 plate (Axion Biosystems, M768-tMEA-48B) at a density of 10,000 cells/well. First, cells were plated in a small 10 uL droplet and incubated for 1 hour at 37°C to ensure proper cell distribution and adhesion to the electrodes. Next, 300 uL of classic neuronal medium was added per well, and a half-medium change was performed every 2 days. At Day 23 in culture, cell activity was recorded on Maestro Pro (Axiom Biosystems) using AxIS software (version 3.11.1.22) for 5 minutes.

### RNA extraction and sequencing

RNA was isolated from cortical neurons using the RNeasy Mini kit (Qiagen, 74104), according to the manufacturer’s protocol. Briefly, the media was removed, and cells were scraped from the dish in Buffer RLT, homogenized 5 times with a 15G needle, passed through the included spin column, and eluted. RNA concentration was quantified via Qubit RNA HS Assay (Invitrogen), and RIN scores were assessed on the BioAnalyzer with the RNA 6000 Nano kit (Agilent, 5067-1511). Samples had an average RIN of 9.15 + 0.65. RNA sequencing was performed by the UAB Genomics Core Lab using 500 ng of RNA as input. Sequencing libraries were made using the NEBNext Ultra II Directional RNA-seq kit (NEB, E7760L) as per the manufacturer’s instructions and using the NEBNext Poly(A) mRNA Magnetic Isolation Module (NEB, E7490). The purified mRNA was fragmented with heat and cations and converted to cDNA with a mixture of random primers for first strand synthesis followed by standard second strand synthesis. The resulting molecules were ligated to a universal adaptor. The Illumina sequences and the unique index information were added via PCR. The resulting cDNA libraries were quantitated using qPCR in a Roche LightCycler 480 with the Kapa Biosystems kit for Illumina library quantitation (Kapa Biosystems, Woburn, MA) prior to cluster generation. The resulting libraries were analyzed on the BioAnalyzer 2100 (Agilent) and quantified by qPCR (Roche). Sequencing was performed on the NovaSeq 6000 (Illumina) with paired-end 100 bp chemistry as per standard protocols. Fastqs were processed using the nf-core/rnaseq (version 3.19.0) pipeline (57) with the --gcBias flag, aligning to the GRCh38.p13 reference, and using the Gencode v47 transcript annotations. DESeq2^16^ was used with default settings to identify differentially expressed genes after filtering out genes with fewer than 10 reads in all three samples within a group. After multiple testing correction using the Benjamini-Hochberg (BH) method^17^, an FDR cutoff of 0.01 and log2 (fold change) > 2 was used to identify differential expression. All analyses were performed using R (R Foundation for Statistical Computing, version 4.3.1). Plots were generated using EnrichedVolcano^18^ and ComplexHeatmap^19^ packages. Gene Ontology (GO) enrichment was performed in R (R Foundation for Statistical Computing, version 4.5.1) using Bioconductor (version 3.21) and topGO packages (version 2.60.1). Gene annotation was provided by org.Hs.eg.db (version 3.21.0). Analyses performed for the Biological Process (BP), Molecular Function (MF), and Cellular Component (CC) ontologies. For each ontology, the enrichment was calculated using Fisher’s exact test as implemented in topGO. GO results were visualized using ggplot2 (version 4.0.0).

### Protein extraction and immunoblotting

Cellular lysates were prepared as described by Gupta et al.^20^ with modifications. The cells were lysed in Pierce™ RIPA Buffer (Thermo Fisher Scientific, 89900), supplemented with Halt™ Protease Inhibitor Cocktail (Fisher Scientific, 78438) and Pierce™ Phosphatase Inhibitor (Thermo Fisher Scientific, A32957) for 2 minutes. Lysates were then scraped, collected into microcentrifuge tubes, and clarified. Protein concentrations for all lysates were determined using the Pierce™ BCA Protein Assay Kit (Thermo Fisher Scientific, 23227). For immunoblotting, 25–30 μg of total protein per sample were separated by SDS-PAGE on Mini-PROTEAN® TGX™ precast gels (Bio-Rad, 4561093) and subsequently transferred to nitrocellulose membranes (Thermo Fisher Scientific, 88018). Membranes were blocked with 5% (w/v) skim milk in Tris-buffered saline containing 0.1% Tween® 20 (TBS-T; Teknova, T9515) for 1 hour at room temperature. Following blocking, membranes were incubated overnight at 4°C with primary antibodies diluted in blocking buffer. The following primary antibodies and dilutions were used: rabbit anti-14-3-3γ (1:10,000, abcam, ab155050), mouse anti-α-Tubulin (1:5,000, Sigma-Aldrich, T5168), mouse anti–β-actin (1:1,000, Thermo Fisher Scientific, MA5-15739), rabbit anti-HSP70 (1:10,000, Proteintech, 10995-1-AP), and rabbit anti-histone H3 (1:1,000 , Thermo Fisher Scientific, PA5-16183). After incubation, membranes were washed three times for 10 minutes each with TBS-T (teknova, T9515) at room temperature with gentle agitation, followed by 1 hour incubation at room temperature with the appropriate horseradish peroxidase (HRP)-conjugated secondary antibodies in the following dilutions: goat anti-mouse IgG (1:10,000, Jackson ImmunoResearch, 115-035-146) and goat anti-rabbit IgG (1:10,000, Jackson ImmunoResearch, 111-035-046). After three additional washes with TBS-T, immunoreactive bands were visualized using the Immobilon® Crescendo Western HRP substrate (Millipore, WBLUR0500). Images were acquired, and band intensities were quantified using ChemiDoc™ MP Imaging System (Bio-Rad, 12003154) and its associated software (version 6.1, Bio-Rad).

### Pharmacology

Neuronal cultures were treated with 10 μM lovastatin (Tocris, 1530) or 10 μM Y27632 (abcam, ab144494) for 1 hour, followed by incubation with Fluo-4 AM dye (Thermo Fisher Scientific, F14201) as described above (a total of 1 hour and 30 minutes of treatment) prior to live cell calcium imaging. Neuronal cultures were treated with 10 μM lovastatin or 10 μM Y-27632 Rho kinase inhibitor for 12 hours prior to RNA and protein isolation, as described above. DMSO (0.1%, v/v) was used as a vehicle control for all conditions.

### Trypsin-induced stress assay

Protein lysates were prepared following the method described by Gupta et al.^20^ with modifications. Neuronal cultures were detached by trypsinization using Trypsin-EDTA (0.05%, Gibco™, 25300054) for 10 minutes. The cell suspension was collected in microcentrifuge tubes and pelleted. After a single wash with phosphate-buffered saline (PBS), the pellet was resuspended directly in standard SDS sample buffer. Protein concentration and immunoblot was performed as described above.

### Data and statistical analysis

All experiments were performed in at least three independent biological replicates (defined as “batches”), in separate rounds of iPSC thawing, plating, culture, neuronal differentiation, and maintenance, unless noted. All statistical analyses were conducted using RStudio version 2024.09.1+394 and R (R Foundation for Statistical Computing, version 4.4.2). To enable batch–aware comparisons, all analyses explicitly tracked biological replicate (“batch”), well, site, and unique ROI identifiers.

Calcium datasets were analyzed as follows: for the calcium baseline analysis, the first frame of each site and ROI was used to quantify intensity levels at the single ROI level for visualization, and the results were analyzed in aggregate at the well level. Baseline fluorescence levels were z-score normalized within each batch to account for batch-to-batch variability. To account for initial fluorescent decay and variability, an additional early-window metric was obtained by averaging frames 1-5 and then used for downstream statistical testing. For analysis of event frequency and amplitude, the fluorescence time series was converted to ΔF/F using a rolling mean baseline. To reduce sensitivity to slow baseline drift, a rolling baseline for each ROI, computed over a fixed window (30 frames), was established, and ΔF/F was calculated. Calcium events were then detected at the ROI level by applying a rolling median and median absolute deviation (MAD). Calcium events were identified as local maxima exceeding a threshold defined as rolling median + 4×MAD. To prevent repeated event counting, we defined a minimum refractory period of 5 frames. Calcium event frequency (in Hz) was defined as the total number of detected events divided by the 30-second recording, and calcium event amplitude was defined as the median ΔF/F value across detected events within each ROI. ROIs with no detected events were retained for frequency analysis but excluded from amplitude analysis.

Although calcium measurements were obtained at single–cell resolution, all reported summary statistics reflect independent biological replicates (defined as separate rounds of iPSC plating, differentiation, and imaging). Single–cell values were first averaged within batch, and batch–level means were then used to compute mean ± SEM across biological replicates (n = 3) for reporting. All statistical testing was performed using linear mixed–effects models (LME) implemented in the lme4 and lmerTest packages in R. Models included genotype, treatment, and their interaction as fixed effects, with batch and batch:well specified as random intercepts to account for hierarchical experimental structure. Individual LME models were fitted for baseline calcium, calcium event frequency, and calcium event amplitude. Statistical inference focused on fixed–effect p–values derived using Satterthwaite’s approximation for degrees of freedom. Interaction terms were tested to determine whether treatment effects differed by genotype, and when genotype interactions were not significant, significant main treatment effects were interpreted as common across genotypes.

Apoptosis analysis was performed by quantifying the intensity of cleaved caspase–3 immunofluorescence at the single–cell level using the built-in MetaXpress software (Molecular Devices). This data was exported at the well level, and analyzed using a linear mixed–effects model with genotype as a fixed effect and batch as a random effect; mean ± SEM values were reported across independent biological replicates (n = 3).

The MAP2 positive area per NeuN–positive neuron was quantified at single–cell resolution using the MetaXpress software (Molecular Devices, LLC) and analyzed with linear mixed–effects models with genotype as a fixed effect and batch as a random effect; mean ± SEM values were reported across independent biological replicates (n = 3).

Western blot band intensities of Trypsin-EDTA-treated samples were normalized to histone H3, to account for cytoskeletal instability and loss of cytoplasmic proteins, using ChemiDoc™ MP Imaging System (12003154) and its associated software (Bio-Rad, version 6.1). Non-trypsinized samples were similarly normalized to β-actin. After normalization, samples were analyzed using paired, batch–level comparisons of log₂ fold–change (lovastatin / DMSO), with statistical significance assessed using one–sample t–tests across independent biological replicates.

### Code availability

All code will be made available on GitHub upon publication.

## RESULTS

### YWHAG^R132C^ iPSC-derived neurons exhibit early cytoskeletal phenotypes

We have characterized a cortical neuron^21^ model of YWHAG Syndrome, differentiated via NGN2 overexpression, using the iPSC iP11NK control line (ALSTEM) edited to carry the R132C mutation (Transcripta Bio) (Figure 1a). Both *YWHAG^+/+^* and *YWHAG^R132C/+^*-derived cortical neurons were positive for the neuronal markers NeuN *(RBFOX3)* and MAP2 (*MAP2*) at d.i.v. 28 (days *in vitro*, Figure 1b). There was a marked reduction in the area of MAP2/NeuN+ neuron (*YWHAG^+/+^* Mean ± S.E.M. = 2857 px^2^ ± 607; *YWHAG^R132C/+^*= 779 px^2^ ± 117; linear mixed effects model (LME) accounting for batch: t(4) = 3.36; p = 0.028, n=3) in *YWHAG^R132C/+^* neurons when compared to isogenic controls (Figure 1c), consistent with the described role of 14-3-3γ as a cytoskeletal scaffolding protein^22^. There was no statistically significant difference in neuronal apoptosis (average intensity of cleaved-caspase 3 per cell *YWHAG^+/+^* Mean ± S.E.M. = 2468 ± 221, *YWHAG^R132C/+^* = 4073 ± 670, LME model, accounting for batch, t(2) = −3.32, p = 0.08, n=3, independent biological replicates, intensity in arbitrary units (Supplementary Figure 1).

**Figure 1:**
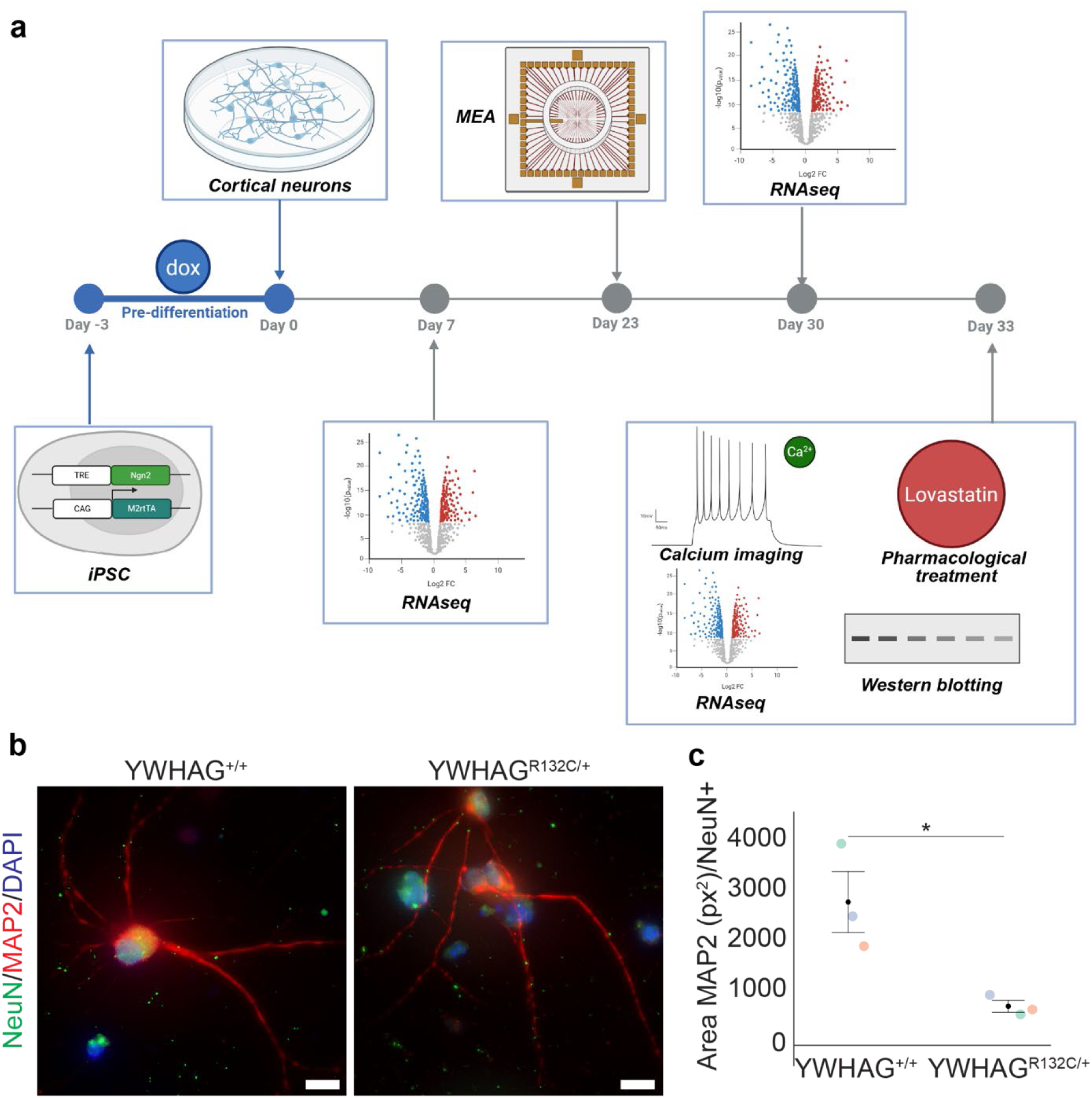
*YWHAG^R132C/+^* characterization strategy and cytoskeletal phenotype. **a)** Strategy for NGN2-induction, differentiation, and characterization of *YWHAG^R132C/+^* cortical neurons. **b)** MAP2 (red) and NeuN (green) labeling of NGN2-derived *YWHAG^R132C/+^*showing reduced neuronal branching compared to isogenic controls. All nuclei were marked with DAPI. **c)** Quantification of total MAP2 area of NeuN+ cells in *YWHAG^R132C/+^* and controls (*YWHAG^+/+^* Mean ± S.E.M. = 2857 px^2^ ± 607; *YWHAG^R132C/+^* = 779 px^2^ ± 117; linear mixed effects model (LME) accounting for batch: t(4) = 3.36; p = 0.028, n=3 independent biological replicates). Colors indicate batches, error bars indicate Mean ± S.E.M.

### YWHAG^R132C/+^ cortical neurons are characterized by a higher calcium baseline, fewer calcium spikes, and reduced network activity

Changes in neuronal excitability are a core component of the epileptic phenotype^23^. Patients with YWHAG Syndrome develop early-onset seizures that eventually lead to developmental abnormalities^2–4^. Little is known about the specific alterations in neuronal excitability associated with 14-3-3 dysfunction, but changes in calcium homeostasis, which modulate neuronal excitability, are frequently observed in developmental epilepsies^24^. To assess the functional phenotypes of differentiated *YWHAG^R132C/+^* neurons, we performed calcium imaging using the Fluo-4 AM calcium indicator (Figure 2a). The *YWHAG^R132C/+^*neurons exhibited a consistently elevated calcium baseline, evident as early as d.i.v. 18 in culture and maintained through d.i.v. 72 (Figure 2a, b; n=3 independent biological replicates, one batch per time point, quantified in Figure 5). The number of calcium spikes at the single-cell level was undetectable (d.i.v. 18, 33) or reduced (d.i.v. 72) in *YWHAG^R132C/+^* neurons (Figure 2c) compared to the isogenic control.

**Figure 2:**
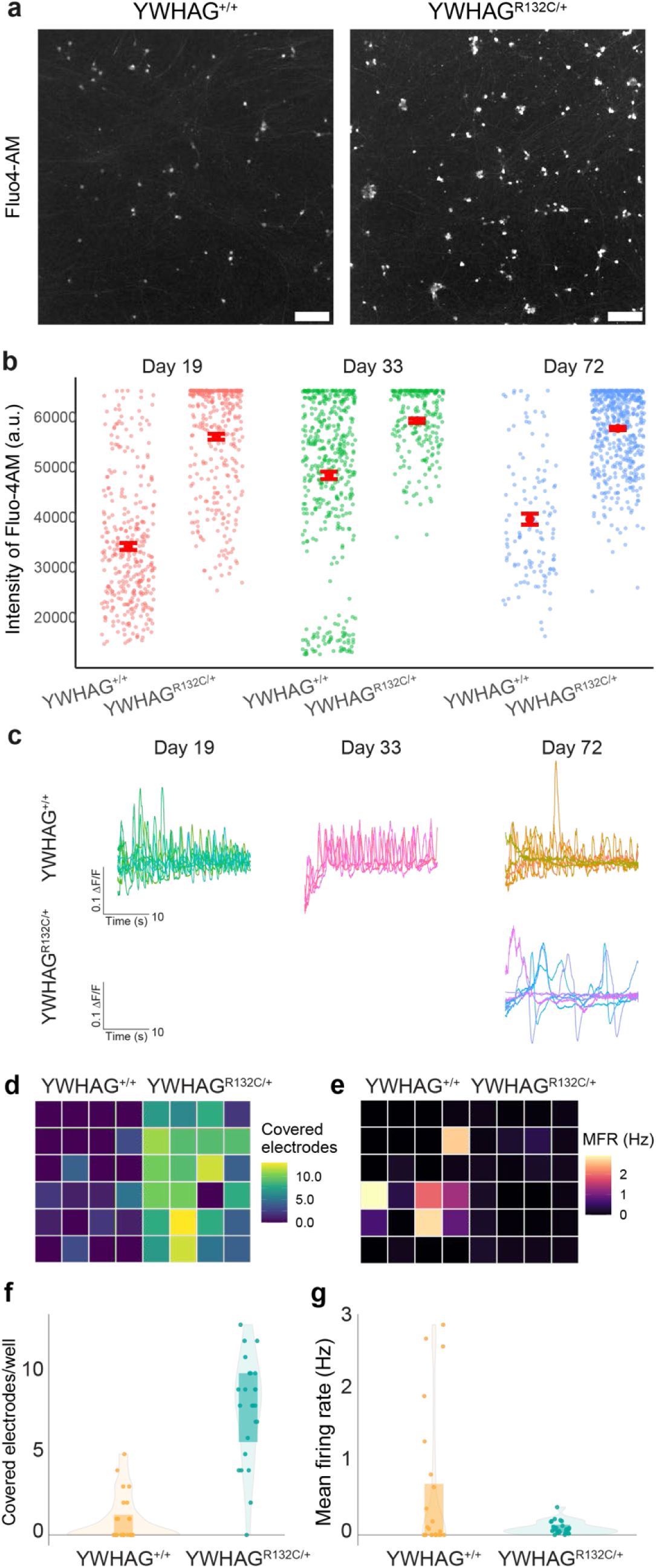
*YWHAG^R132C^* leads to an increase in calcium baseline and a reduction in calcium spikes and network activity. **a)** Calcium baseline staining with Fluo-4 AM. **b)** Quantification of the baseline intensity of calcium labeling with Fluo-4 AM across three independent biological replicates at distinct times in culture. Full analysis and statistical inference are reserved for fully replicated conditions in Figure 5. **c)** Example traces of calcium spikes from **b)** showing reduced or undetectable activity early in *YWHAG^R132C/+^* neuronal cultures. **d-g)** Example MEA data of a single representative batch at d.i.v. 23 (24 wells/condition), demonstrating increased coverage but reduced network activity in *YWHAG^R132C/+^* neurons (n=1).

To validate the effect of alterations to both the calcium homeostasis and the number of spikes in *YWHAG^R132C/+^* neurons at the neuronal network level, we performed a proof-of-concept experiment of multielectrode array (MEA) recordings (n=1, 24 wells/genotype). The *YWHAG^R132C/+^* neurons exhibited increased adhesion to the MEA electrodes, as indicated by broader well coverage (Figure 2d, f), consistent with the described migration defects in *in vivo* models of *YWHAG* overexpression. *YWHAG^R132C/+^* neurons showed reduced network activity and a lower mean firing rate compared to isogenic control neurons (Figure 2e, g).

### YWHAG^R132C/+^cortical neurons have widespread alterations to the transcriptome

14-3-3γ is a binding partner of multiple cytoplasmic and cytoskeletal proteins, potentially modulating complex interactions between downstream pathways^25^. To identify the global effects of *YWHAG^R132C/+^* on the transcriptome of iPSC-derived cortical neurons, we sequenced bulk RNA (RNAseq) at two distinct timepoints: early (d.i.v. 7) and matured (d.i.v. 30), when calcium spikes are consistently detected. Bulk RNAseq of neurons at d.i.v. 7 in culture revealed 2,490 differentially expressed genes (DEGs) upregulated (log2(FC) > 2) and 1,801 DEGs downregulated (log_2_(FC) < 2), indicating widespread alterations to the transcriptome (Figure 3a, b; n=3 biological replicates). At d.i.v. 30, the total number of DEGs was decreased, with 628 genes upregulated and 1628 genes downregulated (n=3 biological replicates) (Figure 3c, d).

**Figure 3:**
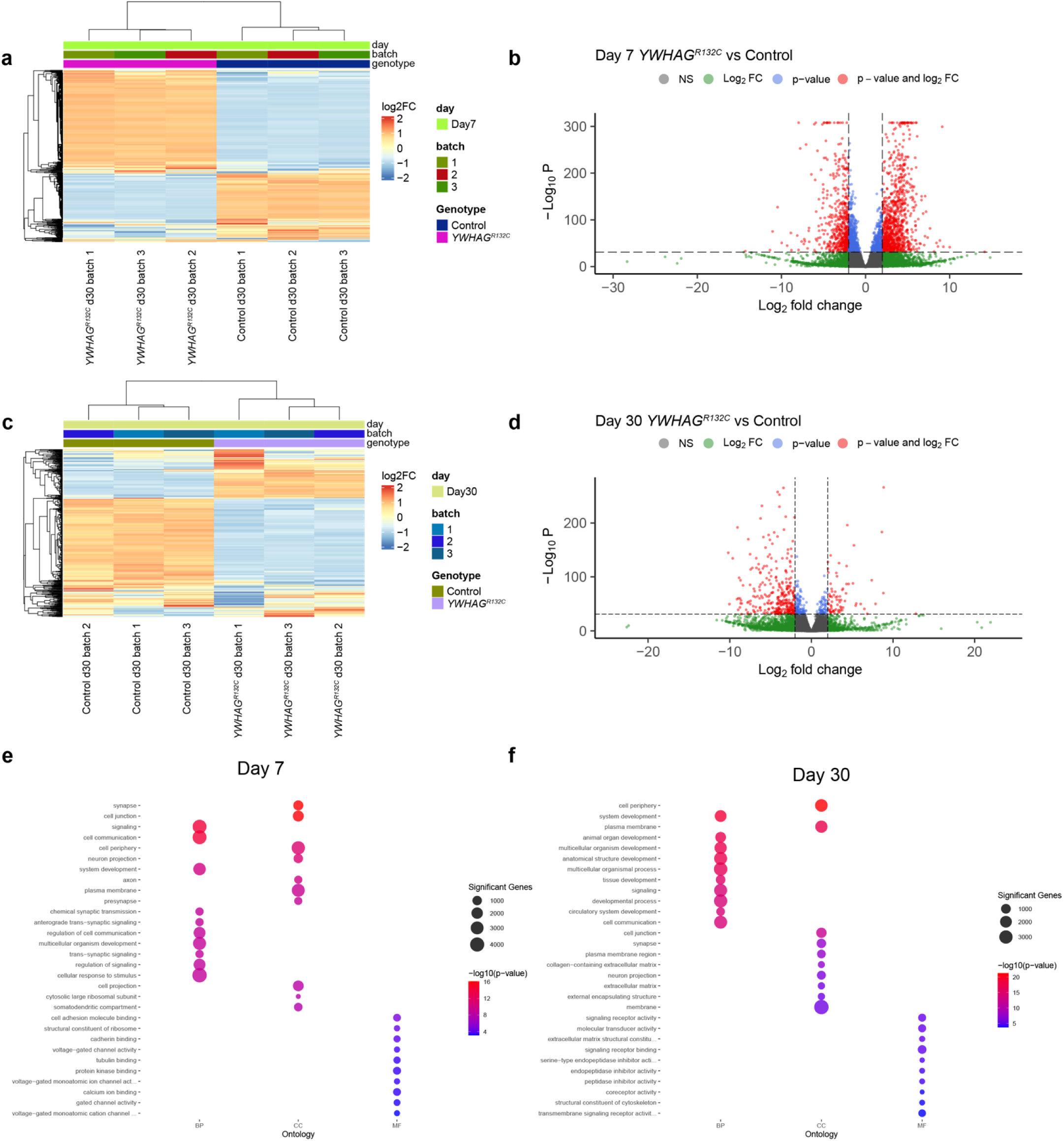
Transcriptomic analysis of *YWHAG^R132C/+^* neurons at 7 and 30 d.i.v. **a, c)** Heatmaps showing differentially expressed genes between *YWHAG^R132C/+^* and isogenic control neurons at d.i.v. 7 (**a**) and d.i.v. 30 (**c**) of differentiation. Expression values are shown as normalized counts scaled per gene across samples. Samples cluster by genotype at both timepoints. **b, d)** Volcano plots of differential gene expression comparing *YWHAG^R132C/+^* and control neurons at d.i.v. 7 (**b**) and d.i.v. 30 (**d**), showing early widespread misregulation that is stabilized with time in culture. Genes are colored according to significance and log_2_(FC), as indicated. **e, f)** Gene Ontology (GO) enrichment analysis of significantly upregulated (red) and downregulated (blue) genes in *YWHAG^R132C/+^* neurons at d.i.v. 7 (**e**) and d.i.v. 30 (**f**). Dot size represents the number of genes contributing to each GO term, and color indicates enrichment significance. RNAseq was performed on n = 3 independent biological replicates per genotype per time point. Differential expression analysis was conducted using DESeq2 (Wald test with Benjamini-Hochberg correction). GO enrichment was performed on significantly differentially expressed genes.

The Gene Ontology (GO) analysis of DEGs in *YWHAG^R132C/+^*neurons revealed a coherent cytoskeletal/adhesion and calcium–handling program across time in culture (Figure 3e, f). The DEGs at d.i.v. 7 showed significant depletion of molecular function (MF) terms associated with multiple ion channels, including voltage-gated and gated channel activities (GO:0022832, GO:0022836) and calcium ion binding (GO:0005509). In addition to channel functions, MF terms associated with cell adhesion molecule binding (GO:0050839), cadherin binding (GO:0045296), and tubulin binding (GO:0015631) were also depleted. The GO terms associated with cellular components (CC), like cell junction (GO:0030054), cell periphery (GO:0071944), synapse (GO:0045202), and neuron projections (GO:0043005), and terms associated with biological processes (BP) of signaling (GO:0023052) and cell communication (GO:0007154) were enriched in d.i.v. 7 DEGs, suggesting a significant role of *YWHAG* in neural function and processes at this stage. The GO analysis of d.i.v. 30 DEGs revealed depletion of MF terms connected to signaling receptor binding (GO:0005102), signaling receptor activity (GO:0038023), and, critically, structural constituent of cytoskeleton (GO:0005200). The enriched GOs for d.i.v. 30 CC also included cell periphery (GO:0071944) and cell junction (GO:0030054), overlapping with the top enrichments from neurons at d.i.v. 7, but ontologies of synapse (GO:0045202) and neuron projections (GO:0043005) shifted towards depletion. Finally, the enriched BP GOs at d.i.v. 30 were associated with the general developmental processes, including system development (GO:0048731), tissue development (GO:0009888), and cell communication (GO:0007154).

### The ROCK pathway regulates YWHAG^R132C/+^ neuronal protein stability and calcium phenotypes

The GO enrichment analysis highlighted the misregulation of pathways associated with cytoskeletal and/or cell adhesion phenotypes in *YWHAG^R132C/+^*cortical neurons, coupled with transcriptional alterations to calcium channels and regulators. These ontologies converge on RhoA/ROCK–actomyosin signaling, which integrates junction/ECM mechanics with calcium homeostasis via mechanotransduction^26–30^. Based on current literature, we identified the transcriptomic network associated with the ROCK pathway, including core genes, upstream/downstream effectors, and focal actin module^31^. At d.i.v. 7, there was a coordinated upregulation of the majority of ROCK-associated transcripts in *YWHAG^R132C/+^*across individual ROCK modules, including upstream regulators (RhoGEFs/GAPs), the core RhoA–ROCK1/2, downstream effectors (LIMK–Cofilin; MLC/myosin II), and focal–adhesion/actin scaffolds (integrins, *TLN1/TLN2*, *VCL*, *PXN*), consistent with increased cytoskeletal stiffness. Interestingly, there was a drastic shift in expression patterns at d.i.v. 30 samples, where the vast majority of ROCK-associated genes were downregulated in *YWHAG^R132C/+^*, consistent with an increase in cytoskeletal instability (Figure 4a).

**Figure 4:**
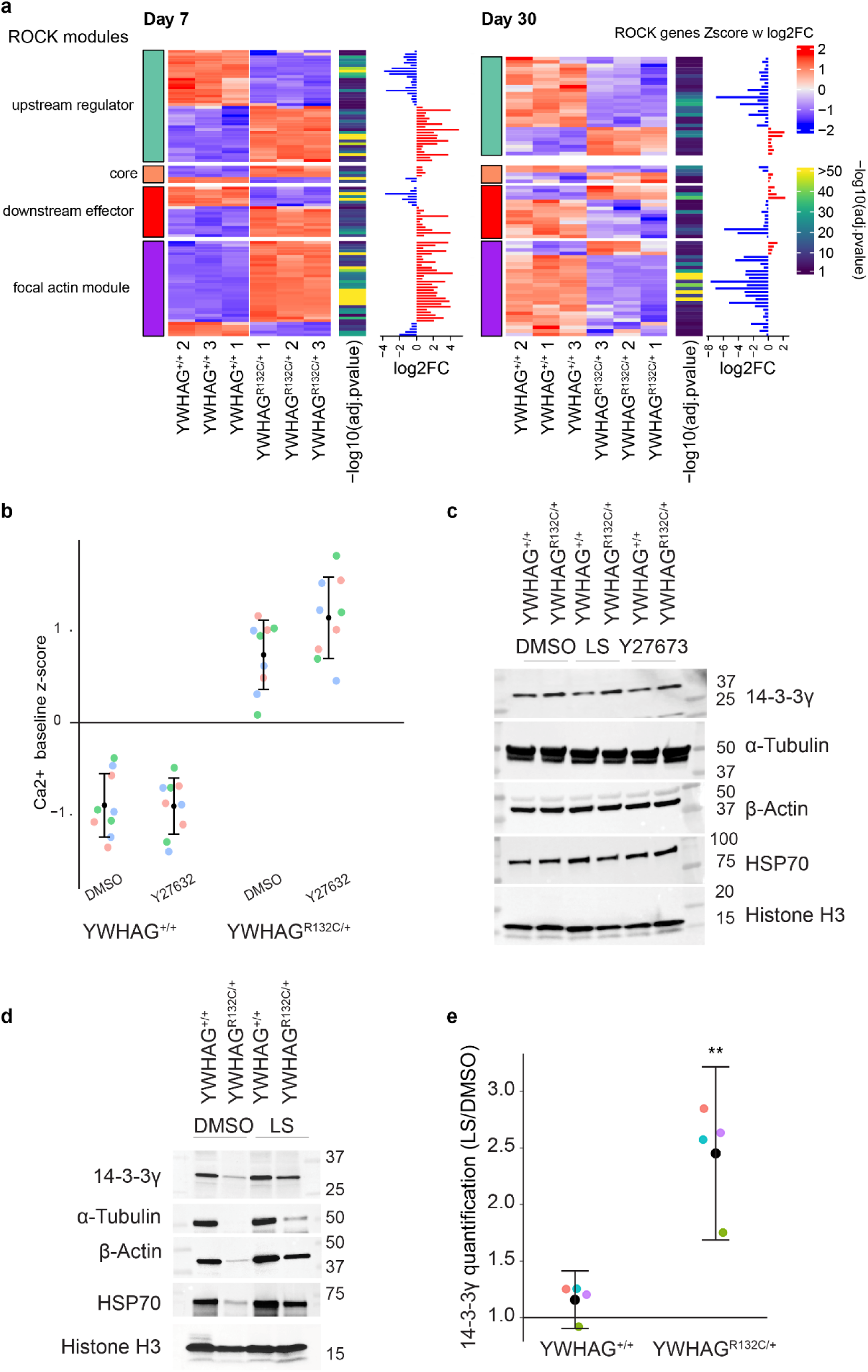
Transcriptomic analysis of ROCK pathway: literature-based core genes, upstream regulators, downstream effectors, and focal actin modules. **a)** Fold change of *YWHAG^R132C/+^*versus control. N=3 independent biological replicates at d.i.v. 7 and d.i.v. 30, matching the RNAseq timepoints in Fig. 3). **b)** Treatment with the ROCK inhibitor Y27632 results in a shift in calcium baseline (stats) **c)** Western blot quantification of protein expression in *YWHAG^R132C/+^*and isogenic control neurons. Modulation of the ROCK pathway with Y27632 and lovastatin did not alter the protein levels of 14-3-3γ. **d)** Trypsin-EDTA (0.05%)-induced cytoskeletal destabilization leads to a drastic loss of cytoskeletal proteins and HSP70. This is partially rescued by lovastatin, which restores protein stability. **e)** Quantification of 14-3-3γ protein levels upon Trypsin-EDTA mediated stress and the lovastatin rescue (*YWHAG^+/+^* Mean FC lovastatin/DMSO ± S.E.M. = 1.16 ± 0.08, *YWHAG^R132C/+^* = 2.45 ± 0.241; one sample t-test on log2FC lovastatin/DMSO; *YWHAG^+/+^* p=0.161; *YWHAG^R132C/+^*p=0.00395; n=4, independent biological replicates, intensity in arbitrary units).

**Figure 5:**
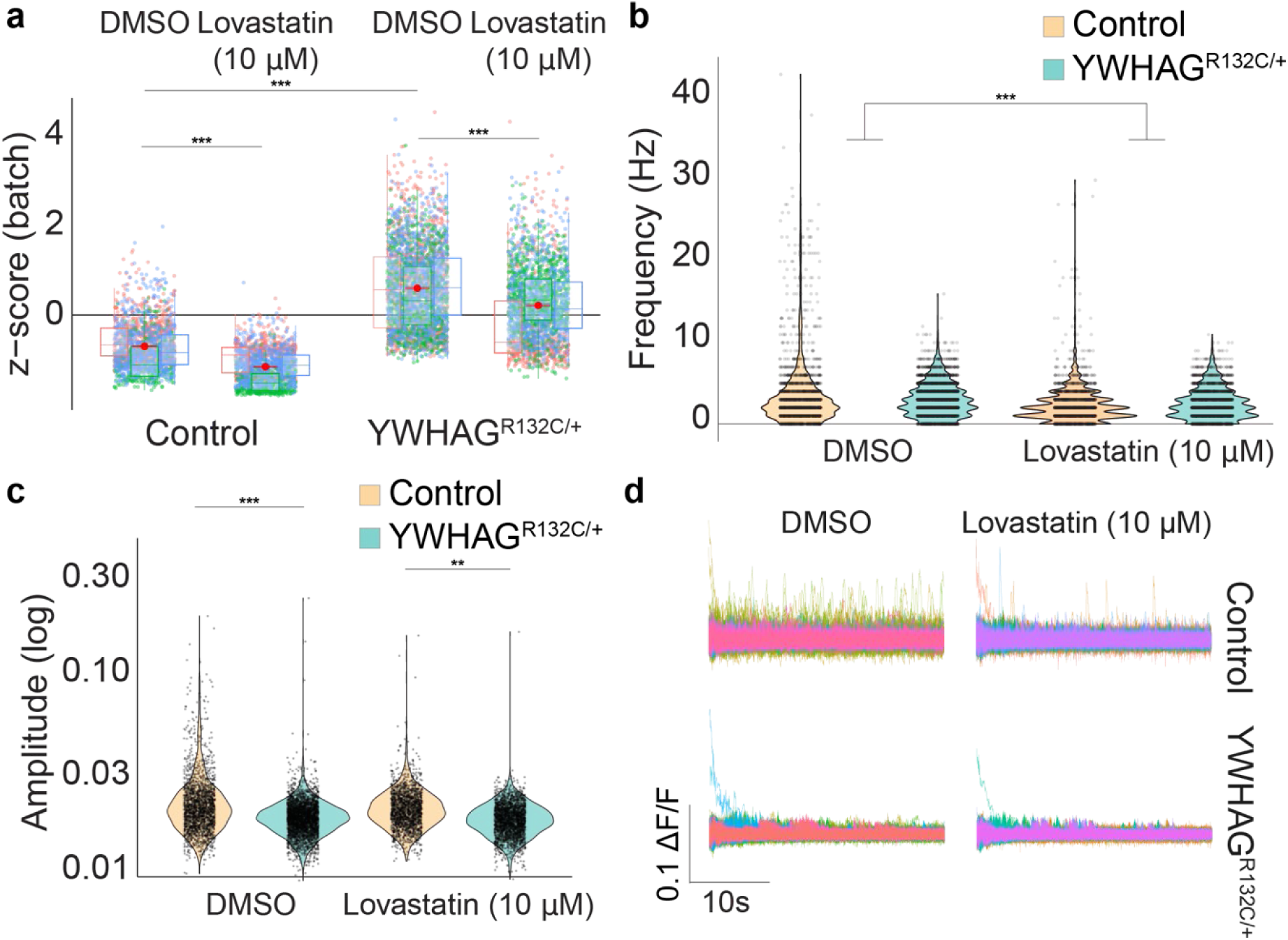
Lovastatin partially rescues the calcium baseline of *YWHAG^R132C/+^* phenotype at 33 d.i.v, but not the amplitude of calcium spikes. a) *YWHAG^R132C/+^* neurons have an elevated calcium baseline when compared to isogenic controls (*YWHAG^+/+^*normalized z-score Mean ± S.E.M = −0.63 ± 0.124, *YWHAG^R132C/+^* = 0.713 ± 0.0730; LME accounting for batch, p<2×10^−16^, n=3 independent biological replicates). Exposure to lovastatin (10 μM) for 1h and 30m was sufficient to lower the calcium baseline, both in *YWHAG^R132C/+^* and control iPSC lines (*YWHAG^+/+^*normalized z-score Mean ± S.E.M = −1.05 ± 0.154, *YWHAG^R132C/+^* = 0.286 ± 0.162; LME accounting for batch, p=0.00412, n=3 independent biological replicates, no significant genotype-treatment interaction). **b)** Frequency (Hz) of calcium spikes was not altered by genotype (*YWHAG^+/+^* Mean frequency ± S.E.M = 3.45 ± 0.485; *YWHAG^R132C/+^* = 3.02 ± 0.248; LME accounting for batch, p=0.09, n=3 independent biological replicates) but lovastatin treatment reduced the frequency in both genotypes (*YWHAG^+/+^* Mean frequency ± S.E.M = 2.36 ± 0.210; *YWHAG^R132C/+^* = 2.45 ± 0.264; LME accounting for batch, p=1.2×10^−4^; n=3 independent biological replicates); and **c, d)** there was a reduction in the amplitude of calcium spikes in *YWHAG^R132C/+^* (*YWHAG^+/+^* Mean amplitude ± S.E.M = 0.0239 ± 0.00170; *YWHAG^R132C/+^* = 0.0201 ± 0.000343; LME accounting for batch, p=3.1×10^−8^; n=3 independent biological replicates), which is not rescued by lovastatin (*YWHAG^R132C/+^* with DMSO = 0.0201 ± 0.000343; *YWHAG^R132C/+^* with lovastatin = 0.0198 ± 0.000511; LME accounting for batch, p=0.98; n=3 independent biological replicates), and the amplitude of *YWHAG^+/+^* calcium spikes remains higher than in *YWHAG^R132C/+^* neurons comparison with (*YWHAG^R132C/+^YWHAG^+/+^* with lovastatin = 0.0224 ± 0.000348, *YWHAG^R132C/+^* with lovastatin = 0.0198 ± 0.000511; LME accounting for batch, p = 0.0018). Colors indicate batches, **d)** sampling of 20 random calcium traces.

To test the functional involvement of the ROCK pathway in *YWHAG^R132C/+^*neuronal phenotypes at d.i.v. 33, we used the competitive ROCK1 and ROCK2 antagonist Y27632, which prevents ROCK function by competing with ATP binding (Figure 4b). ROCK inhibition did not alter calcium homeostasis in *YWHAG^+/+^*neurons after 1 hour and 30 minutes of treatment, but resulted in an upward shift of the calcium baseline in *YWHAG^R132C/+^*, confirming the involvement of ROCK in maintaining calcium homeostasis in *YWHAG^R132C/+^*neurons. Although direct inhibition of ROCK with Y27632 demonstrates pathway involvement, it can rapidly reduce ROCK activity, producing a drastic effect. Therefore, we used lovastatin, which modulates the ROCK pathway through upstream regulation of protein prenylation rather than direct competition, although it also regulates cholesterol biosynthesis, and inhibits L-type calcium channels^32^, and Y27632 to evaluate the effect of ROCK inhibition on neuronal protein expression. Neither Y27632 (10 μM), nor lovastatin (10 μM) after 12 hours of treatment altered protein levels of 14-3-3γ, cytoskeletal proteins α-Tubulin and β-actin, or HSP70 (*YWHAG^R132C/+^* log_2_(FC) lovastatin/DMSO ± S.E.M. = −0.29 ± 0.27, one-sample t-test on log_2_(FC) lovastatin/DMSO, p=0.395, n=3 independent biological replicates).

The reduction in gene expression associated with ROCK-dependent focal actin regulation at d.i.v. 33 (Figure 4a) raised the question of whether cytoskeletal stability, rather than protein synthesis, was affected. To test this, we pre-treated the cells with Trypsin-EDTA (0.05%) immediately before lysis and protein quantification. Remarkably, we saw a drastic reduction in levels of 14-3-3γ, α-Tubulin, β-actin, and HSP70, in *YWHAG^R132C/+^* neurons, suggesting that Trypsin-EDTA digestion of cytoskeletal adhesion molecules leads to a catastrophic failure of cytoskeletal homeostasis but not in the isogenic control (Figure 4d). The same treatment with 10 μM lovastatin for 12 hours was sufficient to partially rescue protein stability, leading to a shift in 14-3-3γ towards control levels and no effect in control neurons (Figure 4e, *YWHAG^+/+^* Mean FC lovastatin/DMSO ± S.E.M. = 1.16 ± 0.08, *YWHAG^R132C/+^* = 2.45 ± 0.241; one sample t-test on log_2_(FC) lovastatin/DMSO; *YWHAG^+/+^* p=0.161; *YWHAG^R132C/+^*p=0.00395; n=4, independent biological replicates, intensity in arbitrary units), accompanied by an parallel increase in α-Tubulin, β-actin, and HSP70. Since lovastatin is associated with the regulation of multiple cellular pathways, we used bulk RNAseq to assess the effects of treatment on ROCK pathway modules. After 12 hours of lovastatin exposure, we observed a robust transcriptional rescue across ROCK modules in *YWHAG^R132C/+^*neurons, with 61/88 ROCK-associated genes shifting directionally towards the control (n=1, Supplementary Figure 2).

### Lovastatin partially rescues YWHAG^R132C/+^elevated calcium baseline but not the calcium spike frequency or amplitude

Lovastatin can modulate intracellular calcium homeostasis indirectly via ROCK or directly by inhibiting L-type calcium channels^32^. We tested whether lovastatin (10 μM) could rescue the calcium homeostasis phenotype in *YWHAG^R132C/+^* neurons at 33 d.i.v. within a 1-hour and 30-minute window of lovastatin treatment (Figure 5). We found that *YWHAG^R132C/+^* neurons had an elevated calcium baseline when compared to isogenic controls (*YWHAG^+/+^*normalized z-score Mean ± S.E.M = −0.63 ± 0.124, *YWHAG^R132C/+^* = 0.713 ± 0.0730; LME accounting for batch, p<2×10^−16^, n=3 biological replicates), and that lovastatin significantly reduced baseline calcium levels in both control and *YWHAG^R132C/+^* groups (*YWHAG^+/+^*normalized z-score Mean ± S.E.M = −1.05 ± 0.154, *YWHAG^R132C/+^* = 0.286 ± 0.162; LME accounting for batch, p= 0.00412, n=3, no significant genotype-treatment interaction), resulting in a partial rescue (Figure 5a). We did not detect a significant difference in the frequency of calcium spikes between *YWHAG^R132C/+^* and *YWHAG^+/+^*neurons (*YWHAG^+/+^* Mean frequency ± S.E.M = 3.45 ± 0.485; *YWHAG^R132C/+^* = 3.02 ± 0.248; LME accounting for batch, p = 0.09, n=3 biological replicates), but lovastatin treatment reduced the frequency in both genotypes (*YWHAG^+/+^* Mean frequency ± S.E.M = 2.36 ± 0.210; *YWHAG^R132C/+^* = 2.45 ± 0.264; LME accounting for batch, p = 1.2×10^−4^; n=3 biological replicates; Figure 5b, d). The *YWHAG^R132C/+^* neurons were also characterized by reduced calcium spike amplitude (*YWHAG^+/+^* Mean amplitude ± S.E.M = 0.0239 ± 0.00170; *YWHAG^R132C/+^* = 0.0201 ± 0.000343; LME accounting for batch, p = 3.1×10^−8^; n=3 biological replicates). Lovastatin did not rescue calcium amplitude in *YWHAG^R132C/+^* neurons (*YWHAG^R132C/+^* with DMSO = 0.0201 ± 0.000343; *YWHAG^R132C/+^* with lovastatin = 0.0198 ± 0.000511; LME accounting for batch, p=0.98), and *YWHAG^R132C/+^* neurons maintain lower calcium amplitudes despite the rescue of baseline calcium levels (*YWHAG^+/+^* with lovastatin = 0.0224 ± 0.000348, *YWHAG^R132C/+^* with lovastatin = 0.0198 ± 0.000511; LME accounting for batch, p = 0.0018, n=3 biological replicates, Figure 5c, d). Together, these results support a role of the ROCK pathway in the cytoskeletal and calcium phenotype of *YWHAG^R132C/+^*cortical neurons and suggest a decoupling between maintenance of calcium homeostasis and the frequency and amplitude of calcium events.

## DISCUSSION

YWHAG Syndrome poses a challenge for disease modeling due to the limited availability of clinical samples and the small patient population. Further, the role of the 14-3-3 family, including 14-3-3γ, as scaffolding proteins during cytoskeletal assembly and cellular homeostasis makes it challenging to parse the effects of *YWHAG* mutations on neuronal function. In this study, to better understand the downstream effects of the *YWHAG^R132C^* mutation on neuronal function, growth, and maturation, we differentiated iPSC-derived *YWHAG^R132C/+^* cortical neurons and paired isogenic controls. NGN2-induced differentiation of *YWHAG^R132C/+^*revealed maturation-associated phenotypes of cytoskeletal instability, with increased baseline calcium, and decreased event frequency and amplitude. Although most epileptic phenotypes can be associated with hyperexcitability, a higher calcium baseline can lead to network hypoexcitability, resulting in an epileptic phenotype, such as in the case of interneurons in *SCN1A* and loss-of-function (LOF) mutations in Dravet Syndrome^33^, and reduced excitability in cortical excitatory neurons in LOF *SCN2A*^34^.

Transcriptomic analysis of both early and matured *YWHAG^R132C/+^* revealed a biphasic involvement of the ROCK pathway, initially as an enforcer of cytoskeletal rigidity, and later as a stabilizer in a compromised cytoskeletal state. Direct inhibition of the ROCK pathway further shifted the *YWHAG^R132C^*-associated calcium baseline away from the control. However, treatment with lovastatin for 1 hour and 30 minutes was sufficient to partially rescue the calcium baseline phenotype towards control, but did not restore the frequency or amplitude of calcium events. ROCK inhibition for 12 hours did not alter 14-3-3γ, cytoskeletal, or HSP70 protein levels, whereas treatment with lovastatin slightly reduced 14-3-3γ protein levels. Strikingly, Trypsin-mediated perturbation of the cytoskeleton led to a catastrophic loss of 14-3-3γ, cytoskeletal proteins, and HSP70 in *YWHAG^R132C/+^* neurons, a loss that was partially restored after 12 hours of lovastatin treatment, suggesting that lovastatin acts at the level of protein stabilization, reducing compensatory protein synthesis.

Lovastatin is an inhibitor of 3-hydroxy-3-methylglutaryl coenzyme A (HMG-CoA) reductase and is known under the generic names Mevacor and Altoprev^35^. It is an FDA-approved drug for hypercholesterolemia, but it also attracted interest as a potential anti-seizure drug candidate when it was shown to decrease inflammatory markers during the epileptogenesis period in the hippocampal region of epileptic rats^36^, suggesting a potential neuroprotective effect after status epilepticus. By inhibiting HMG–CoA reductase, lovastatin reduces prenylation, the synthesis of isoprenoids like farnesyl and geranylgeranyl that are essential for the prenylation of small GTPases, including the Rho family^37–39^. Finally, lovastatin has been shown to inhibit L-type calcium channels in smooth muscle cells, directly modulating calcium homeostasis^32^. Lovastatin’s role in multiple cellular pathways beyond cholesterol synthesis and blood-brain barrier permeability has expanded its potential use beyond hypercholesterolemia to neurological and neurodegenerative diseases, including Fragile X syndrome and Parkinson’s disease^40,41^. Of particular note, in mouse models of Fragile X syndrome, lovastatin had a beneficial effect while Simvastatin did not show a marked effect^42^. We found that lovastatin modulates *YWHAG^R132C^* phenotypes, particularly calcium homeostasis, where it decouples the calcium baseline from calcium spike frequency and amplitude, and cytoskeletal stability. This modulation of both calcium and cytoskeletal phenotypes by lovastatin suggests potential druggable nodes that can be targeted for YWHAG Syndrome and broader conditions associated with YWHAG dysfunction.

The broader effects of 14-3-3γ-dependent regulation of neural function are evident in conditions such as Down Syndrome (DS) and Williams Syndrome, in which direct involvement of 14-3-3γ has been established. Individuals with fetal DS exhibit significantly downregulated 14-3-3γ expression in the cortex^43^, whereas aged DS patients show increased protein abundance in several brain regions, including the cortex ^44^. Williams Syndrome, a neurodevelopmental condition affecting multiple organs^45^, presents atypically when the deletion of chromosome 7q11.23 includes *YWHAG*^46^. Patients carrying the *YWHAG* deletion suffer from epilepsy, which is not a symptom in the typical clinical presentation^47^. Strikingly, there is also strong evidence of the 14-3-3γ engagement in neurodegenerative disorders like Alzheimer’s and Parkinson’s. In mouse models, 14-3-3γ appears to promote α-synuclein aggregation^48^ and co-localizes with Lewy bodies^49^; remarkably, there is an indirect link whereby lovastatin can alleviate α-synuclein aggregation^50^. In Alzheimer’s disease patients, the expression of the YWHAG protein is increased in the cortex of aged patients^44^, while it also co-localizes with neurofibrillary tangles in the hippocampal region^51^. Nonetheless, the precise role of 14-3-3γ in those progressive conditions remains to be elucidated.

In conclusion, we have described an isogenic iPSC-derived cellular model of YWHAG Syndrome, allowing for the study of functional and molecular outcomes associated with 14-3-3γ mutations. We have found that *YWHAG^R132C/+^* neurons have disrupted cytoskeletal homeostasis, which is maintained via ROCK. Inhibition of ROCK leads to a critical homeostatic failure, and this perturbed state can be stabilized by lovastatin. Further, stabilization with lovastatin decouples the calcium baseline from calcium event frequency and amplitude, indicating a *YWHAG*-associated, cytoskeletal-dependent, reconfiguration of calcium homeostasis. Together, these results establish testable molecular pathways and therapeutic targets for YWHAG Syndrome and YWHAG-associated disorders.

## CONTRIBUTIONS

AMS, AG, designed and performed experiments and statistical analysis. AT and RDS executed experiments. DRB and LR performed RNA isolation and bulk RNA-sequencing analysis. JP designed experiments and performed statistical analysis.

## ACKNOWLEDGEMENTS

We would like to thank the YWHAG Foundation for their encouragement and financial support. The UAB Genomics Core Lab is supported by the NIH S10 mechanism 1S10OD032422-01.

---

**Supplementary Figure 1:**
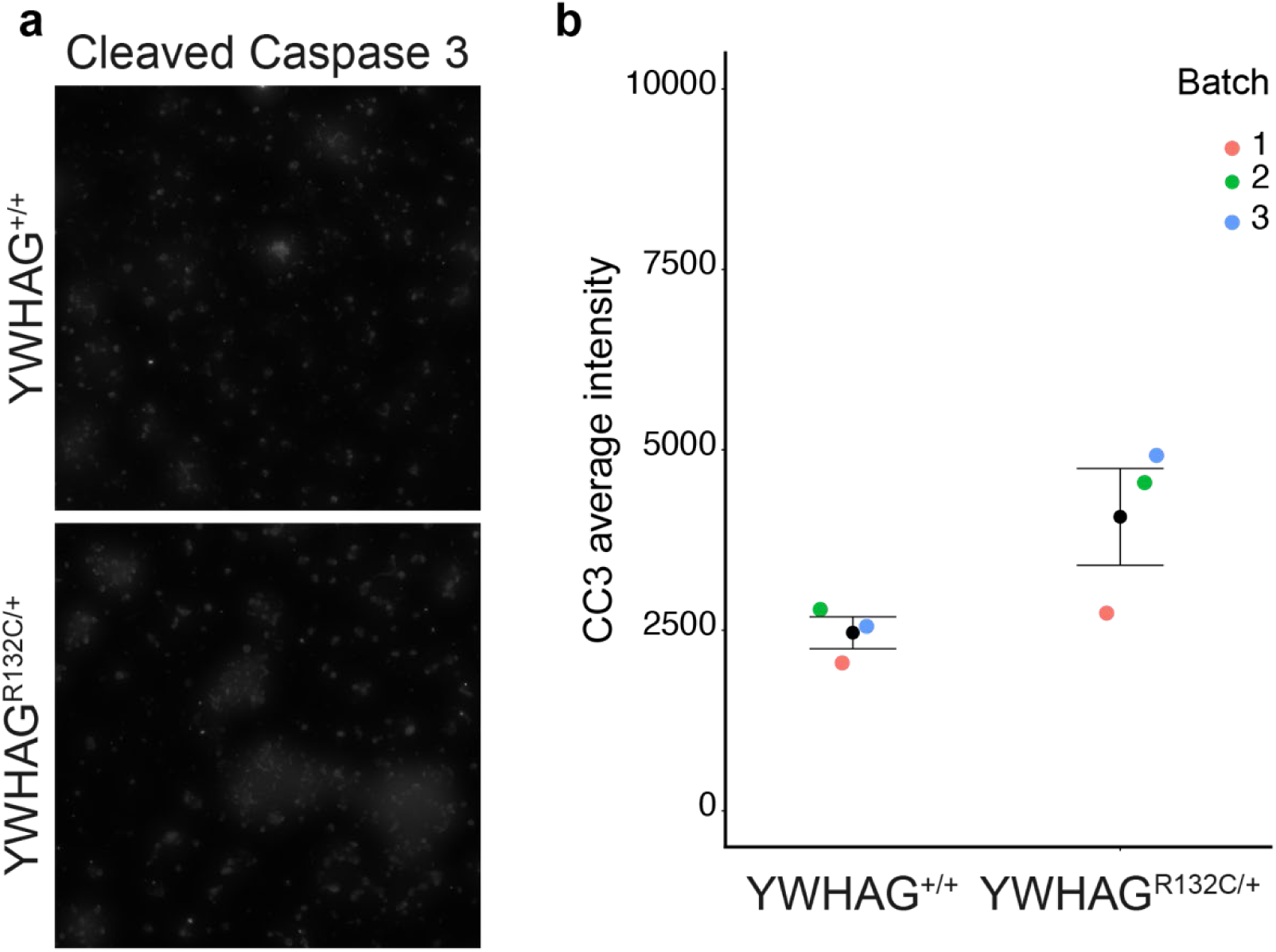
Quantification of the pro-apoptotic marker Cleaved Caspase-3 by immunostaining. **a)** Cleaved Caspase-3 (CC3) staining in *YWHAG^R132C/+^* and control neurons. **b)** Quantification of total CC3 average intensity in *YWHAG^R132C/+^*and controls (average intensity of cleaved-caspase 3 per cell *YWHAG^+/+^* Mean ± S.E.M. = 2468 ± 221, *YWHAG^R132C/+^*= 4073 ± 670, LME model, accounting for batch, t(2) = −3.32, p = 0.08, n=3, independent biological replicates, intensity in arbitrary units). Colors indicate batches, error bars indicate Mean ± S.E.M.

**Supplementary Figure 2:**
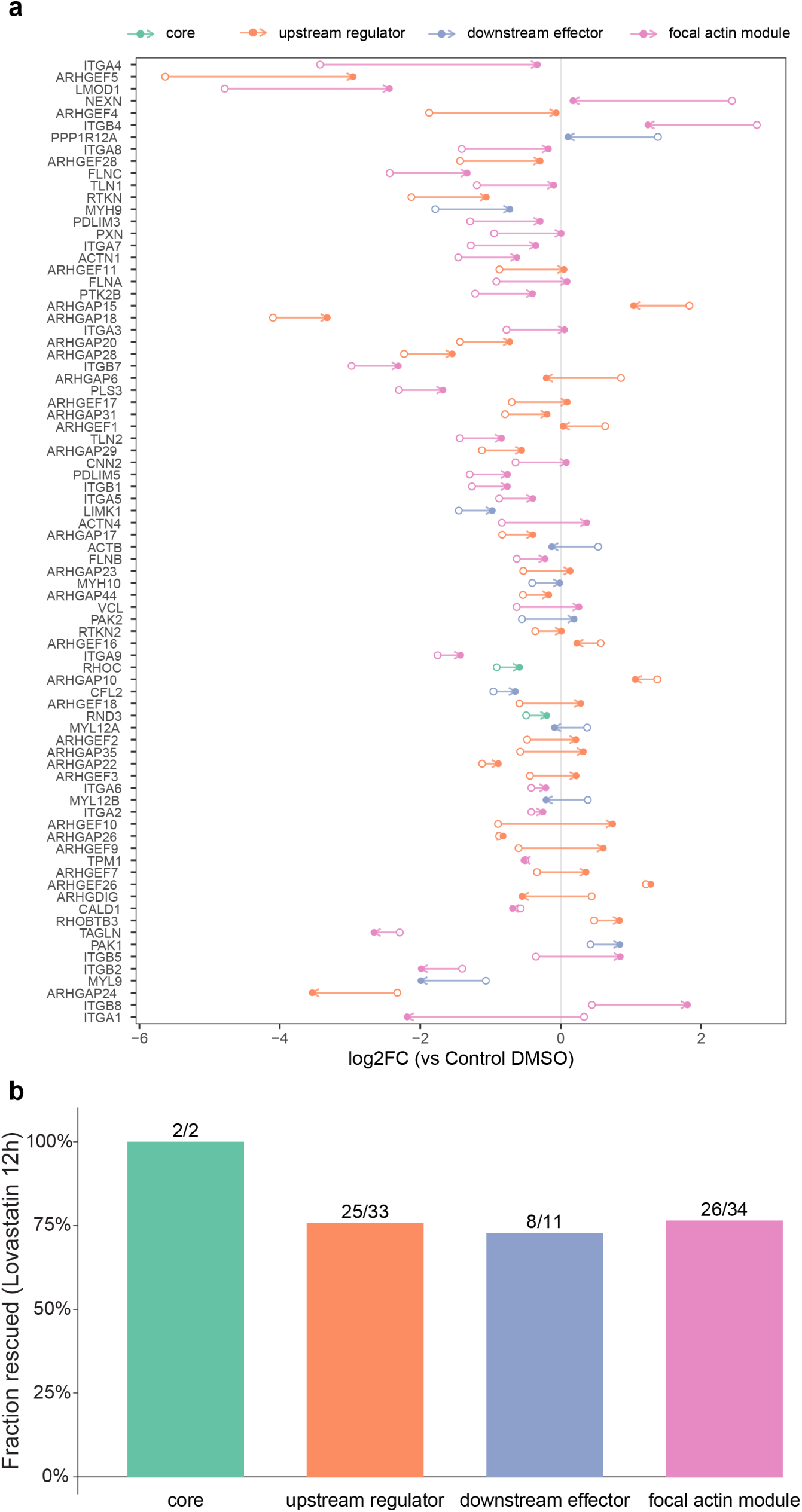
Rescue of ROCK pathway-related genes after 12 hours of lovastatin (10 μM) treatment. Included genes had |log2FC| > 0.3, and a rescue was defined as a reduction in magnitude toward control levels with |log2FC| > 0.2. n=1.

